# Evidence for a widespread third system for bacterial polysaccharide export across the outer membrane comprising a composite OPX/β-barrel translocon

**DOI:** 10.1101/2022.04.04.486985

**Authors:** Johannes Schwabe, María Pérez-Burgos, Marco Herfurth, Timo Glatter, Lotte Søgaard-Andersen

## Abstract

In Gram-negative bacteria, secreted polysaccharides have multiple critical functions. In Wzx/Wzy- and ABC transporter-dependent pathways, an outer membrane (OM) polysaccharide export (OPX) type translocon exports the polysaccharide across the OM. The paradigm OPX protein Wza_*E. coli*_ is an octamer, in which the eight C-terminal domains form an α-helical OM pore, and the eight copies of the three N-terminal domains (D1-D3) a periplasmic cavity. In synthase-dependent pathways, the OM translocon is a 16- to 18- stranded β-barrel protein. In *Myxococcus xanthus*, the secreted polysaccharide EPS is synthesized in a Wzx/Wzy-dependent pathway. Here, using experiments and computational structural biology, we characterize EpsX as an OM 18-stranded β-barrel protein important for EPS synthesis and identify AlgE, a β-barrel translocon of a synthase-dependent pathway, as its closest structural homolog. We also find that EpsY, the OPX protein of the EPS pathway, only consists of the periplasmic D1 and D2 domains and lacks the domain for spanning the OM (henceforth ^D1D2^OPX protein). *In vivo*, EpsX and EpsY mutually stabilize each other, supporting their direct interaction. Based on these observations, we propose a model whereby EpsY and EpsX make up a novel type of translocon for polysaccharide export across the OM. Specifically, in this composite translocon, EpsX functions as the OM-spanning translocon together with the periplasmic ^D1D2^OPX protein EpsY. Based on computational genomics, similar composite systems are present widespread in Gram-negative bacteria. This model provides a framework for these proteins’ future experimental characterization.

## Introduction

Most bacteria secrete one or more polysaccharides. These polysaccharides protect against environmental stresses and phage infection, contribute to surface colonization and biofilm formation, have important functions in beneficial and pathogenic human-, animal- and plant-microbe interactions, provide the basis for serotyping and several anti-bacterial vaccines, and have many applications in the food, pharmaceutical and medical industry ^1-3^. Here, we focus on export of polysaccharides across the outer membrane (OM) in Gram-negative bacteria.

Secreted polysaccharides are large, chemically diverse molecules. Their synthesis hinges on three mechanisms and their export across the OM in Gram-negative bacteria on two known mechanisms ^4,5^. In Wzx/Wzy-dependent pathways, biosynthesis is initiated on the cytoplasmic side of the IM by a phosphoglycosyltransferase (PGT). Subsequently, glycosyltransferases (GTs) add monosaccharides to generate the repeat unit. The Wzx flippase “flips” individual repeat units across the IM to the periplasm, where the Wzy polymerase polymerizes them. On the periplasmic side, the polysaccharide co-polymerase (PCP), an integral IM protein with an extended periplasmic domain, regulates polymerization and polysaccharide transfer across the periplasm to the OM ^5,6^. A protein of the OM polysaccharide export (OPX) family exports the polysaccharide across the OM ^7,8^. Wza of *Escherichia coli* is the best-studied OPX protein. In contrast to other pore-forming OM proteins, the octameric Wza spans the OM using an α-helical barrel ^7^. The periplasmic part of the OPX protein interacts with the oligomeric PCP to establish a conduit for the polysaccharide to reach the OM ^6,8-10^. In ABC transporter-dependent pathways, synthesis of the polysaccharide is also initiated on the cytoplasmic side of the IM and fully completed before translocation across the IM by an ABC transporter ^5^. Transfer across the periplasm involves a PCP, and the polysaccharide is exported across the OM by an OPX protein ^5,6^. Synthase-dependent pathways differ considerably from these two pathways and consist of only three core components ^4^. An IM-embedded synthase with GT activity synthesizes the polysaccharide and, in parallel, facilitates its translocation across the IM. A tetratricopeptide repeat (TPR)-containing protein contributes to polysaccharide transfer across the periplasm. The polysaccharide is exported across the OM through an integral OM 16- to 18-stranded β- barrel protein as described in structural work on PgaA and BcsC of *E. coli*, and AlgE of *Pseudomonas aeruginosa* ^11-13^.

Here, we combine experiments, computational structural biology and genomics to provide evidence supporting a novel type of polysaccharide OM export mechanism widespread in Gram-negative bacteria. In the suggested system, a short OPX protein that only comprises periplasmic domains and lacks the domain for spanning the OM functions together with an OM β-barrel protein to generate a composite OPX/β-barrel translocon.

## Results & Discussion

### Myxobacterial gene clusters for secreted polysaccharides encode an OM β-barrel protein

The Gram-negative deltaproteobacterium *Myxococcus xanthus* secretes three polysaccharides, i.e. exopolysaccharide (EPS), spore coat polysaccharide (SPS) and biosurfactant polysaccharide (BPS), using three dedicated Wzx/Wzy-dependent pathways (Figure S1) ^14^. In all three systems, the gene annotated as encoding the OPX protein (EpsY, ExoA, WzaB) is syntenic with a gene encoding a protein of unknown function (EpsX, ExoB, MXAN_1916). This synteny is largely conserved in the orthologous myxobacterial gene clusters (Figure S1).

EpsX, ExoB and MXAN_1916 have a type 1 signal peptide based on sequence analysis. AlphaFold structural models (Methods) predict with high confidence that they fold into 18-stranded antiparallel β-barrels with an elliptical shape and a central channel (Figure 1A, Figure S2A-C). As expected, the parts of the proteins predicted with low confidence correspond to extracellular loops connecting the antiparallel β–strands (Figure S2A-C). The three AlphaFold models could readily be superimposed (Figure S2D), documenting that the proteins have the same fold overall. ExoB is important for SPS biosynthesis by an unknown mechanism ^15^, and MXAN_1916 is an OM protein ^16^, but it is not known whether it is important for BPS synthesis. Altogether, these data support that EpsX, ExoB and WzaB are integral OM 18-stranded β-barrel proteins. To investigate the function of these proteins and their orthologs, we focused on EpsX.

**Figure 1.**
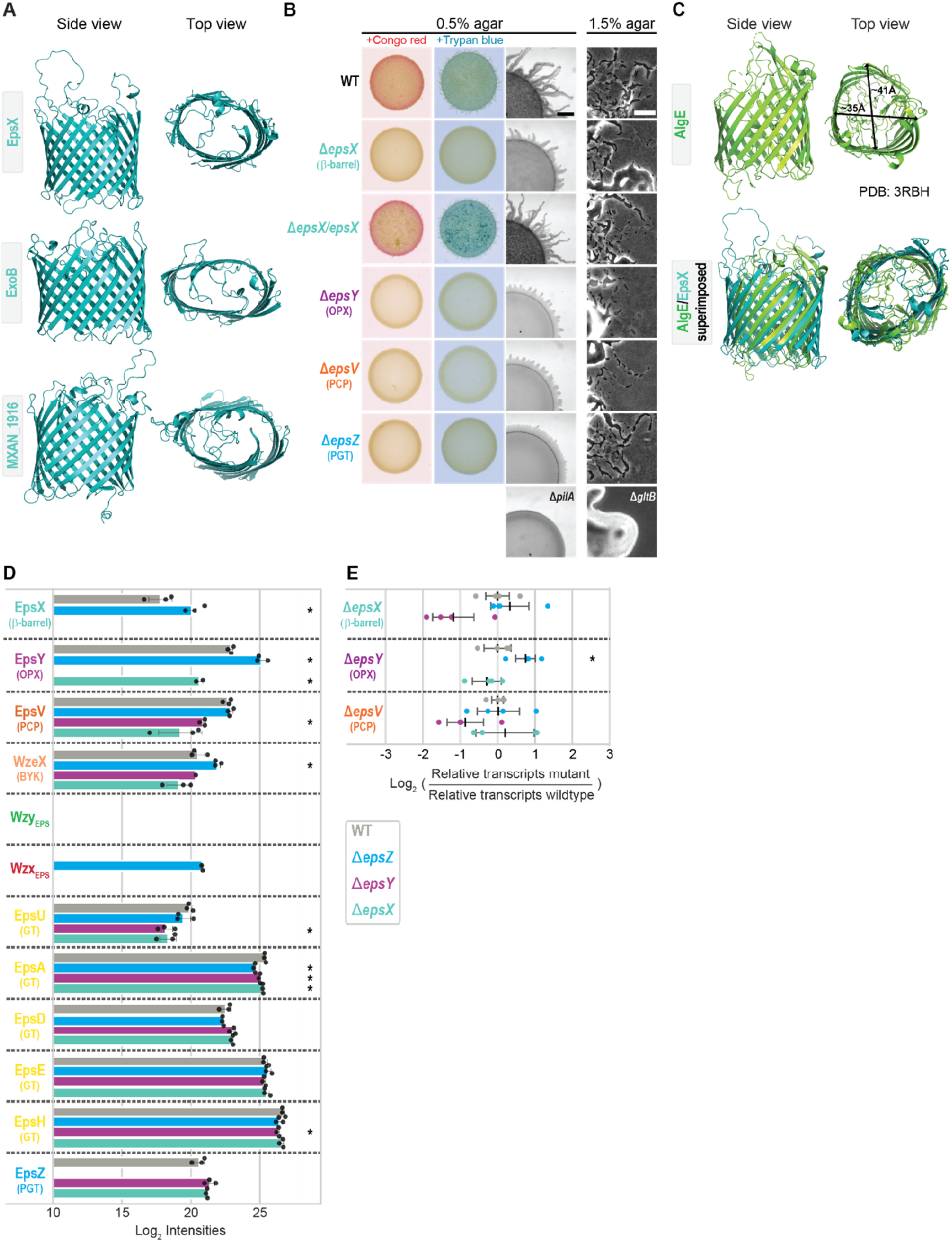
The 18-stranded β-barrel protein EpsX is an integral part of the EPS pathway. **A**. AlphaFold models of EpsX, ExoB and MXAN_1916. Proteins are oriented based on the N- and C-terminus of OM β-barrel proteins being periplasmic ^58^. Model ranks 1 are shown. **B**. Phenotypic characterization of Δ*epsX* mutant. Two left columns, cells were placed on 0.5% agar supplemented with 0.5% CTT and Congo red or Trypan blue and incubated for 24h. Two right columns, T4P-dependent and gliding motility were tested on 0.5% and 1.5% agar, respectively, supplemented with 0.5% CTT and images recorded after 24h. The Δ*pilA* mutant, which lacks the major pilin of the T4P ^59^, and Δ*gltB* mutant, which lacks a component of the gliding motility machinery ^22^, served as negative controls for T4P-dependent and gliding motility, respectively. In the complementation strain, *epsX* was expressed from the *pilA* promoter on a plasmid integrated in a single copy at the Mx8 *attB* site. Scale bars: 1mm and 50μm (left, right). Similar results were obtained in three independent experiments. **C**. Comparison of AlgE and EpsX. Upper panel, lateral and top views of the solved structure of AlgE (PDB 3RBH) ^11^. Arrows indicate the external diameter of the β-barrel. Lower panel, superimposition of the solved structure of AlgE and the EpsX AlphaFold model. EpsX is colored in teal. EpsX aligns to AlgE with an RMSD of 6.035Å over 1501 C_*α*_. **D**. EpsX and EpsY mutually stabilize each other and EpsY stabilizes EpsV. Protein amounts in whole-cell proteomes of *M. xanthus* strains were determined using LFQ mass spectrometry-based proteomics (Methods). Normalized Log_2_ intensities of Eps proteins in the indicated strains are shown. Missing bars indicate that the proteins were not detected. Data points represent three biological replicates. Error bars, standard deviation (STDEV) based on these replicates. * *P*<0.05 Welch’s test. WzeX is important for EPS synthesis and was proposed to act as the BY kinase partner of EpsV ^14,18^. **E**. RT-qPCR analysis of *epsV, epsY* and *epsX* transcripts levels. Total RNA was isolated from cells grown as in **D**. Data are shown as Log_2_ transcripts in a mutant relative to WT. Individual data points represent four biological replicates with each two technical replicates and are colored according to the strain analyzed. Center marker and error bars represent mean and STDEV. * *P*<0.05 Welch’s test. Source data for **A, D, E** are provided in the Source Data file.

### EpsX is important for EPS biosynthesis

To investigate the function of EpsX, we generated an in-frame deletion in *epsX* (Δ*epsX*). Using plate-based colorimetric assays with Congo red or Trypan blue as readouts of EPS synthesis, we observed that wild-type (WT) synthesized EPS. The Δ*epsX* mutation, similarly to the Δ*epsZ*, Δ*epsV* and Δ*epsY* mutations that inactivate genes for proteins in the EPS pathway (Figure S1) ^17-19^, caused strongly reduced EPS synthesis (Figure 1B). EPS is important for type IV pili (T4P)-dependent motility ^14^. Consistently, while WT formed colonies with long flares at the edge characteristic of T4P-dependent motility, and the Δ*pilA* negative control formed smooth-edged colonies, the Δ*epsX* mutant generated short flares at the colony edge as previously observed for the Δ*epsZ*, Δ*epsV* and Δ*epsY* mutants ^17-19^ (Figure 1B). The Δ*epsX* mutant, similarly to WT and the Δ*epsZ*, Δ*epsV* and Δ*epsY* mutants, displayed the single cells at the colony edge characteristic of gliding motility, while the Δ*gltB* negative control did not (Figure 1B). The EPS biosynthesis and motility defects of the Δ*epsX* mutant were complemented by ectopic expression of *epsX* (Figure 1B). We conclude that EpsX is important for EPS synthesis and likely an integral component of the EPS pathway.

In the *E. coli* and *Klebsiella pneumoniae* Wzx/Wzy-dependent pathways for capsule biosynthesis, the OM 18-stranded β-barrel protein Wzi is important for cell surface-anchoring of the capsule but neither for its biosynthesis nor its export ^20,21^ arguing that Wzi and EpsX have different functions. Moreover, while the β-barrel in the solved structure of Wzi is circular ^21^ (Figure S2E), the EpsX β-barrel in the AlphaFold model is elliptical (Figure 1A). Wzi also contains an N-terminal α-helical bundle that occludes the periplasmic side of the β-barrel, and extracellular loops that fold into and occlude the β-barrel on the extracellular side ^21^. By contrast, EpsX lacks the N-terminal α-helical bundle, and the extracellular loops, although modeled with low confidence (Figure S2A), do not fold into the β-barrel, indicating that EpsX is open to the periplasm and possibly also to the cell exterior. This structural comparison also supports the conclusion that EpsX has a function different from Wzi.

Subsequently, by searching for structural homologs of EpsX using Foldseek (Methods), we identified the OM translocon AlgE of the synthase-dependent pathway for alginate export in *P. aeruginosa* ^11^ as the closest structural homolog. Similar to EpsX in the AlphaFold model, the 18-stranded β-barrel of AlgE in the solved structure has an elliptical shape (Figure 1C). The two proteins could readily be superimposed except for extracellular loops in AlgE that fold into the β-barrel and partially occlude the pore (Figure 1C).

### EpsX and EpsY mutually stabilize each other

There are several reported examples in *M. xanthus* of interacting OM, periplasmic and IM proteins that stabilize each other ^22-25^. Consequently, to identify potential EpsX interaction partners, we performed whole-cell label-free quantitative (LFQ) mass spectrometry-based proteomics (Methods) focusing on the Eps proteins.

In WT, we detected all Eps proteins except for the integral IM proteins Wzx_EPS_ and Wzy_EPS_ (Figure 1D). In the Δ*epsX* mutant, the OPX protein EpsY accumulated at a significantly reduced level (Figure 1D). Conversely, in the Δ*epsY* mutant, EpsX was not detected, and accumulation of the PCP EpsV was significantly reduced (Figure 1D). By contrast, the Δ*epsZ* mutant, which lacks the PGT for initiating EPS synthesis, had slightly decreased (EpsA), increased (Wzx_EPS_, EpsY and EpsX), or WT levels of the Eps proteins (Figure 1D). These observations support that the decreased accumulation of EpsY in the Δ*epsX* mutant and EpsX and EpsV in the Δ*epsY* mutant is not caused by lack of EPS biosynthesis and export *per se*. Some GTs are also slightly but significantly reduced in the Δ*epsX* and Δ*epsY* mutants (Figure 1D). Here, we focused on the significant accumulation dependency of EpsX, EpsY and EpsV. Importantly, reverse-transcriptase quantitative PCR (RT-qPCR) provided evidence that the changes in EpsX, EpsY and EpsV levels are independent of transcription in the Δ*epsX* and Δ*epsY* mutants (Figure 1D). We conclude that the OM β-barrel proein EpsX and the OPX protein EpsY mutually stabilize each other and EpsY stabilizes the PCP EpsV.

These findings agree with OPX and PCP proteins interacting in the periplasm ^8-10^. They also support the proposal that EpsX, EpsY and EpsV form a complex that spans the cell envelope. To investigate how these proteins could interact, we first analyzed EpsY and its homologs.

### Myxobacterial OPX proteins are short and only comprise two periplasmic domains

In the Wza_*E. coli*_ octamer, individual protomers have four structural domains (D1-D4) ^7^ (Figure 2A): The N-terminal D1-D3 are periplasmic, with D1 containing the characteristic polysaccharide export sequence (PES) motif. D2 and D3 are structurally related and have a β-grasp fold ^26,27^. The C-terminal D4 is an amphipathic α-helix, which is inserted in the OM. In the octamer, the eight copies of D1-D3 generate a periplasmic cavity, and the eight C-terminal α-helices create the α-helical barrel in the OM (Figure 2A). Wza has a type 2 signal peptide and is N-terminally acylated ^7^, which is important for OM integration ^26,28^.

**Figure 2.**
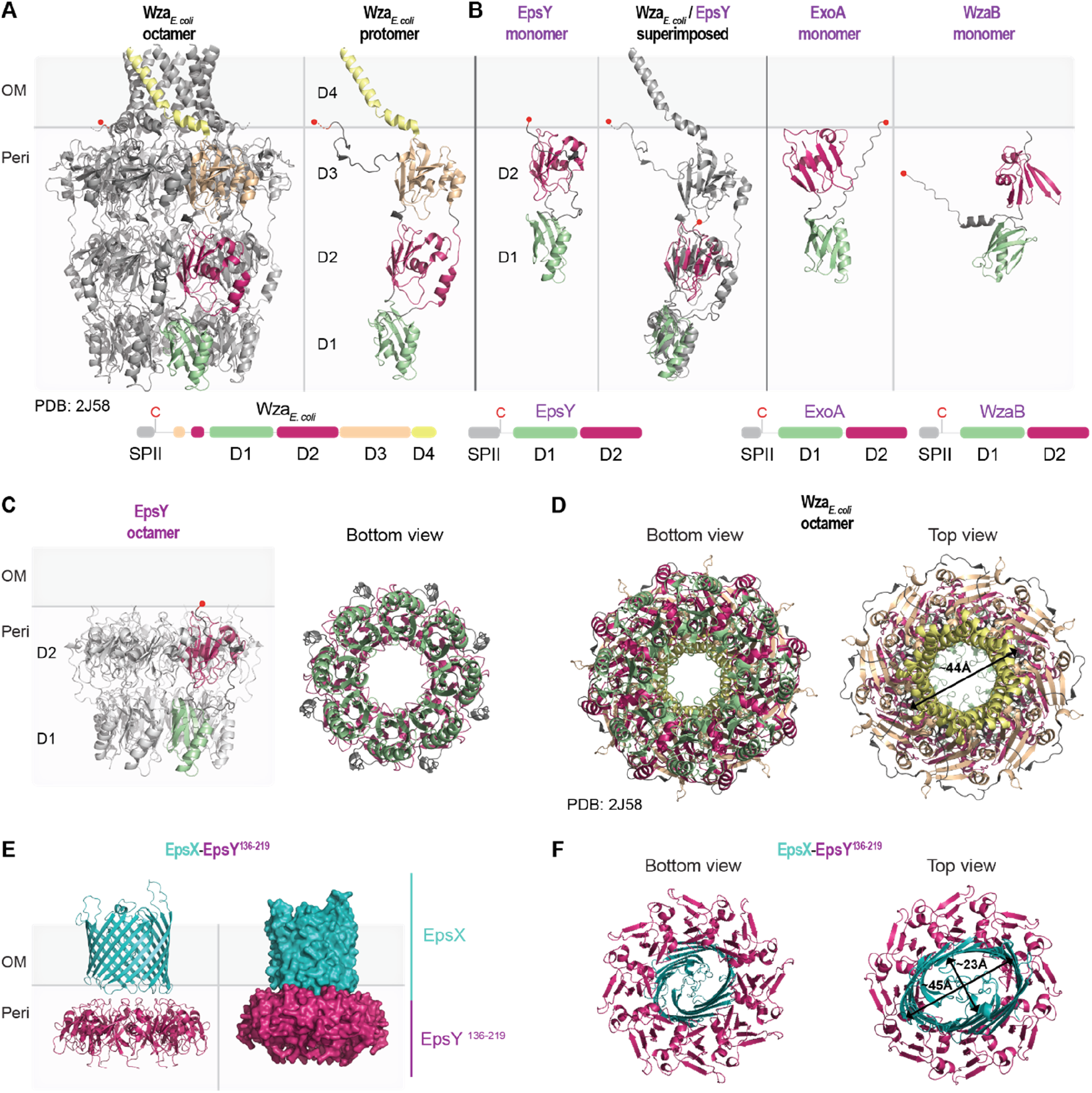
Structural characterization of the EpsY ^D1D2^OPX protein alone and in complex with EpsX, its partner 18-stranded β-barrel protein. **A**. Structure of Wza_*E. coli*_. Left panel, the solved structure of octameric Wza (PDB 2J58) ^7^. Right panel, individual Wza protomer. The four domains of Wza are labeled D1-D4. Light green, D1; Dark pink, D2; Light orange, D3; Yellow, D4. The acylated N-terminal cysteine is indicated with a red ball and placed at the inner leaflet of the OM. Lower panel, domain organization of Wza. **B**. AlphaFold model of EpsY. Left panel, lateral view of EpsY monomer as predicted by AlphaFold. Right panel, superimposition of a Wza protomer from the solved structure (grey) and the EpsY model. The EpsY monomer aligns to the Wza protomer with an RMSD of 3.306Å over 904 C_*α*._ Right panels, AlphaFold models of ExoA and WzaB monomers. In all three AlphaFold models, the two domains are labeled D1 and D2 and colored according the homologous domains in Wza. The acylated N-terminal cysteine is indicated with a red ball; note that the acylated N-terminal cysteine of WzaB is not modeled “on top” of D2, but the confidence in the relative position of this residue is low (Figure S3C). Model rank 1 is shown for all structures. Lower panels, domain organization of EpsY, ExoA and WzaB. SPII, type 2 signal peptide. **C**. AlphaFold-Multimer model of octameric EpsY. Left panel, one protomer is colored as in **B**. Right panel, bottom view of octameric EpsY with all protomers colored as in **B**. Model rank 1 is shown. **D**. Structure of Wza. All eight protomers are colored as in **A**. Arrow indicates the external diameter of the α-helical pore. In the bottom view, the tyrosine residues that form the so-called tyrosine ring are present in the loops extending into the central channel. **E, F**. AlphaFold-Multimer model of a heterocomplex of octameric EpsY^136-219^ and an EpsX monomer. In the heterocomplex, the EpsY^136-219^ octamer is colored as D2 in **B** and EpsX is colored teal. In **E**, the right panel is a surface-rendered representation. Model rank 1 is shown. In **F**, arrows indicate the diameter of the β-barrel. Model rank 1 is shown. Source data for **B, C, E** are provided in the Source Data file.

Based on sequence analysis and a high confidence AlphaFold structural model, EpsY only contains D1 with the PES motif and D2 and lacks D3 and, most strikingly, D4 (Figure 2B; Figure S3A). The two EpsY domains could readily be superimposed on the corresponding domains in a Wza protomer (Figure 2B). Using Alphafold-Multimer (Methods), EpsY with high confidence could generate an octamer in which D1 and D2 form a stacked, ring-like structure with a central cavity, similar to the Wza counterparts (Figure 2C; cf. Figure 2A, D; Figure S4A). The Wza octamer is closed at the periplasmic base of D1 by a so-called tyrosine ring (Figure 2D); however, this tyrosine ring is lacking in the EpsY octamer (Figure 2C). As in the monomer model (Figure 2B), the acylated N-terminal cysteine would be placed on top of D2 (Figure 2C), facilitating the OM association of octameric EpsY. From here on, we refer to OPX proteins that only comprise D1 and D2 as ^D1D2^OPX proteins.

Remarkably, EpsY orthologs in other myxobacteria are also of the ^D1D2^OPX type (Figure S5). In addition, the OPX proteins for SPS (ExoA) and BPS (WzaB) export, as well as their orthologs, are of the ^D1D2^OPX type (Figure 2B; Figure S3B-C; Figure S5). Except for one, all these ^D1D2^OPX proteins have a type 2 signal peptide and are predicted lipoproteins (Figure S5). Thus, all 48 identified OPX proteins in myxobacteria are ^D1D2^OPX proteins and, most strikingly, lack D4 for generating the OM α-helical pore (Figure S5). Consistent with these findings, a previous classification of OPX proteins ^6^ defined a subgroup consisting of short OPX proteins that included EpsY and ExoA.

### A ^D1D2^OPX protein and an 18-stranded β-barrel protein may form a translocon for OM polysaccharide export

^D1D2^OPX proteins lack the domain for spanning the OM and, generally, co-occur with 18-stranded β-barrel proteins (Figure S1, Figure S5). Moreover, the 18-stranded β-barrel proteins are structurally similar to the OM translocon AlgE (Figure 1C), indicating that they can support polysaccharide export across the OM. The ^D1D2^OPX protein EpsY and the 18-stranded β-barrel protein EpsX mutually stabilize each other, supporting that they interact. Based on these three lines of evidence, we hypothesized that a ^D1D2^OPX protein functions together with a β-barrel protein to create a composite ^D1D2^OPX/β-barrel translocon.

To test this hypothesis, we intended to generate a heterocomplex consisting of eight EpsY molecules and one EpsX molecule using AlpfaFold-Multimer. However, this amount of sequence information is computationally highly demanding to analyze. Instead, we, therefore, generated a heterocomplex consisting of EpsY D2 (aa 136-219), which is predicted to be close to the OM (Figure 2B-C), and full-length EpsX. In a high confidence Alphafold-Multimer structural model, EpsX is placed on top of D2 in the EpsY^136-219^ octamer (Figure 2E, Figure S4B) in an arrangement strikingly similar to that of the α-helical barrel “on top” of D3 in the Wza octamer (Figure 2A). Moreover, the diameters of the α-helical barrel in Wza and the EpsX β-barrel in the EpsX/EpsY^136-219^ heterocomplex are similar (Figure 2D, F).

To test the specificity of the AlphaFold-Multimer prediction of the EpsX/EpsY^136-219^ complex, we attempted to generate a model of an AlgE/EpsY^136-219^ heterocomplex. Even in the best of the five predicted models, the AlgE β-barrel clashes with the ring structure of EpsY^136-219^, and the confidence in the relative positioning of AlgE to octameric EpsY^136-219^ is low (Figure S4C). Thus, despite EpsX and AlgE being close structural homologs, the low confidence of the AlgE/EpsY^136-219^ model suggests high specificity in the EpsX/EpsY^136-219^ heterocomplex, as expected for a functionally relevant protein complex.

Altogether, this computational approach and the mutually dependent stability of EpsX/EpsY support that these two proteins form a complex. In this complex, EpsX spans the OM, and EpsY is periplasmic and associated with the OM via the N-terminal lipid group and the interaction to EpsX.

### PCPs in *M. xanthus* may have an extended periplasmic domain

The IM PCP proteins interact with their cognate OPX proteins in the periplasm using a region rich in α-helices ^8-10,29^. In the case of the Wzc-Wza_*E. coli*_ complex, the PCP Wzc_*E. coli*_ in the solved structure extends ∼60Å from the IM into the periplasm ^29^ (Figure 3A), while the periplasmic part of Wza extends ∼100Å from the OM into the periplasm, with each ring contributing ∼30Å ^7^. This gives an estimated periplasmic height of a Wzc-Wza complex of ∼160Å.

**Figure 3.**
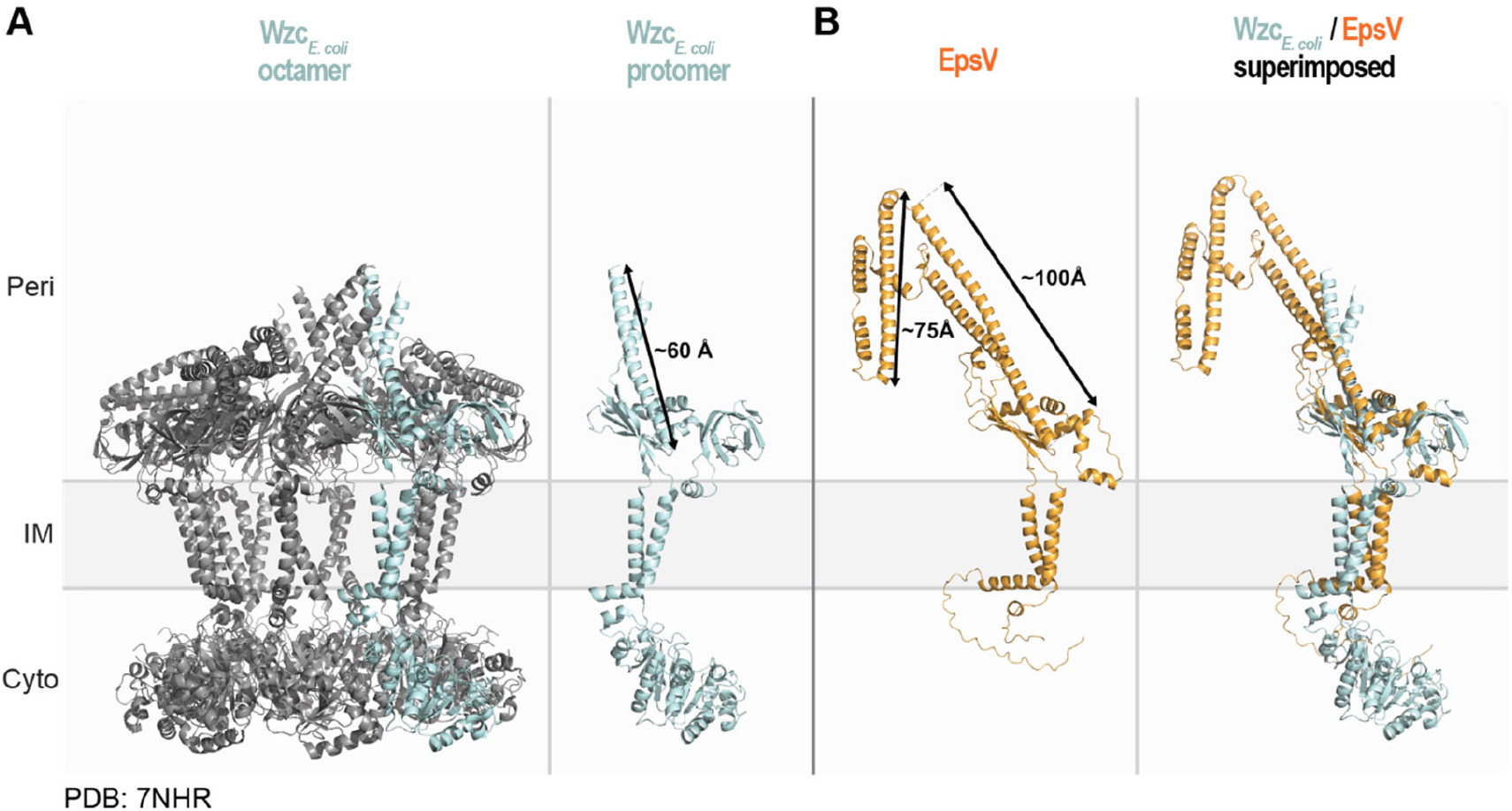
Structural characterization of the PCP EpsV. **A**. The solved structure of octameric Wzc_*E. coli*_ (PDB 7NHR) ^29^. Left panel, the protein is colored in grey and one protomer in light blue. Right panel, individual Wzc protomer in class 2 conformation ^29^. Note, that individual protomers have different conformations in the octamer. Arrow indicates the length of the extended α-helical stretch. **B**. AlphaFold model of EpsV. Arrows indicate the length of the α-helical stretches. Model rank 1 is shown. Right panel, superimposition of a protomer from the solved structure of Wzc and the EpsV model. EpsV aligns to Wzc with an RMSD of 4.610Å over 727 C_*α*._ Source data for **B** is provided in the Source Data file.

EpsY is important for the stability of the PCP EpsV (Figure 1D), supporting that the two proteins interact. This raises the question of how the EpsY ^D1D2^OPX protein would be able to bridge the periplasm together with its EpsV PCP partner. A high confidence AlphaFold model of monomeric EpsV supports that it has two IM trans-membrane α-helices and a periplasmic domain rich in α-helices (Figure 3B; Figure S6A), similar to other PCPs ^6^. The periplasmic part of EpsV is composed of two α-helical stretches with a length of ∼100Å and ∼75Å and connected by linkers. Depending on the conformation of the α-helical stretches in EpsV, the EpsY/EpsV complex could, thus, have a height of ∼160-240Å across the periplasm, supporting that they can jointly span the periplasm.

Similarly, monomeric ExoC of the SPS and WzcB of the BPS pathways have long α-helical periplasmic regions in high confidence AlphaFold models (Figure 3B; Figure S6B-C). Altogether, these *in silico* analyses support the idea that ^D1D2^OPX proteins function together with a PCP with an extended periplasmic part.

### Coupled genes for short periplasmic OPX proteins and OM β-barrel proteins are widespread in Gram-negative bacteria

41 of the 48 myxobacterial genes for ^D1D2^OPX proteins are syntenic with a gene encoding an 18-stranded β-barrel protein (Figure S1; Figure S5). We took advantage of this observation to assess bioinformatically how widespread short, periplasmic OPX proteins are and whether they are coupled with an OM β-barrel protein. Specifically, we identified OPX candidates in 6,607 fully sequenced prokaryotic genomes across 49 phyla using the PES motif (Pfam 02563) as described ^6^. To substantiate that the identified proteins are part of a polysaccharide biosynthesis pathway, we only included OPX candidate genes with a polysaccharide biosynthesis gene within five genes upstream or downstream (Methods). In total, we identified 4,257 OPX proteins, and subsequently used 2,749 representative proteins to determine whether they have a C-terminal α-helix and/or are coupled to an OM β-barrel protein (Methods). To determine the coupling of OPX proteins and OM β-barrel proteins, we searched for genes encoding OM β-barrel proteins within five genes of a gene encoding an OPX protein (Methods). After verifying the β-barrel fold and OM localization computationally (Methods), we identified 486 such OM β-barrel proteins.

The 2,749 OPX proteins range in length from 138 to 1,728 aa (Figure 4A). Interestingly, the length distribution of the 486 OPX proteins with a coupled β-barrel protein is highly skewed towards shorter OPX proteins (Figure 4A-B). Based on the size distribution of OPX proteins with a coupled β-barrel protein, and because EpsY, ExoA and WzaB are 217 aa, 190 aa and 204 aa, respectively, we empirically defined a size cut-off for short OPX proteins of 280 aa (Figure 4A-B). We built a phylogenetic tree of all 2,749 OPX proteins based on their PES motif and found these proteins in 28 phyla (Figure 4C). We then located the branches of the tree that correspond to OPX proteins with a size of ≤280 aa, and found that those with a coupled β-barrel protein are present in 10 phyla (Figure 4C).

**Figure 4.**
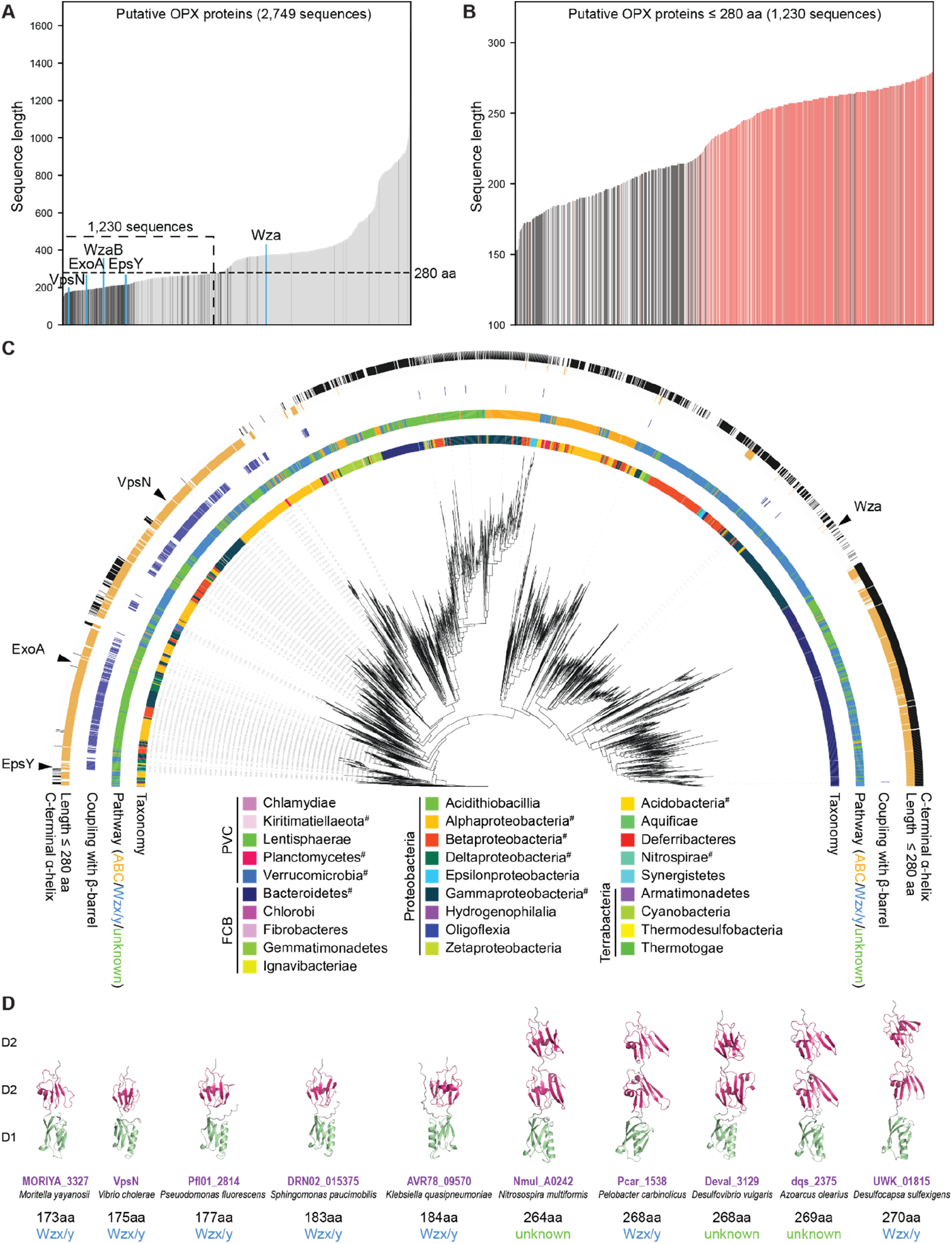
Computational genomics of OPX proteins. **A**. Length distribution of 2,749 OPX proteins. OPX proteins encoded within a distance of five genes of a gene encoding a β-barrel protein are shown in dark grey, and the remaining OPX proteins are in light grey. OPX proteins discussed in the text are highlighted in blue. The 1,230 short OPX proteins are indicated based on the upper size limit of ≤280 aa. **B**. Length distribution of OPX proteins ≤280 aa. 453 OPX proteins with a coupled β-barrel protein and no C-terminal α-helix are in dark grey; 535 OPX proteins with no coupled β-barrel protein but a C-terminal α-helix are in red. Proteins not matching these criteria are in light grey. **C**. Maximum-likelihood phylogenetic tree built from the PES motif of the 2,749 OPX proteins. The inner ring is colored based on NCBI taxonomy (https://www.ncbi.nlm.nih.gov/taxonomy/). PVC, FCB and Terrabacteria are superphyla, the phylum Proteobacteria is divided into classes. # indicates that this phylum contains OPX protein(s) ≤280 aa coupled to a β-barrel protein(s). The second ring indicates the pathway assigned to individual OPX proteins. The third ring indicates in dark blue coupling with a β-barrel protein. The fourth ring indicates in orange OPX protein ≤280 aa. The fifth ring indicates in black whether an OPX protein has a C-terminal α-helix. **D**. AlphaFold models of indicated OPX proteins. Domains are labeled D1 and D2 and colored according to the homologous domains in Wza (Figure 2A). Model rank 1 is shown for all structures. Source data for **A, B, C, D** are provided in the Source Data file.

1,230 OPX proteins are ≤280 aa, and among these, 453 proteins have a coupled β-barrel protein (Figure 4B-C). Conversely, 93% of the β-barrel proteins are coupled to an OPX protein ≤280 aa. Interestingly, among the 777 OPX proteins ≤280 aa and without a coupled β-barrel protein, 535 have a C-terminal α-helix (Figure 4B-C) and are mostly found in the Bacteroidetes (Figure 4C). Thus, generally, we observe a dichotomy among the 1,230 OPX proteins ≤280 aa. Typically, these OPX proteins have either a coupled β-barrel protein and no C-terminal α-helix (453 proteins) or no coupled β-barrel protein but a C-terminal α-helix (535 proteins). For the latter group of OPX proteins, we suggest that they likely function similarly to Wza.

Focusing on the 453 OPX proteins ≤280 aa with a coupled β-barrel protein and no C-terminal α-helix, we found, based on genomic context, that 185 are part of a Wzx/Wzy-dependent pathway, 14 of an ABC transporter-dependent pathway, and 287 were not assigned to a pathway because the closest polysaccharide biosynthesis gene encodes a PCP and some of these proteins can be difficult to assign to a specific type of pathway ^6^. Among the short OPX/β-barrel couples identified, we found VpsN/VpsM of *Vibrio cholerae* (Figure 4A, C-D). These two proteins are important for *Vibrio* polysaccharide (VPS) synthesis, biofilm formation and intestinal colonization in a mouse model ^30^, thus, validating the outlined computational approach. Analysis of the domain architecture of the short OPX proteins is difficult because, except for the PES motif in the D1 domain, their sequences are not well-conserved.

Therefore, to analyze the domain architecture of the 453 short OPX proteins with a coupled β-barrel protein, we analyzed the domain structure of five proteins ≤200 aa and five proteins >250 aa. Intriguingly, the five proteins with a length <200 aa all had the ^D1D2^OPX architecture, while the proteins >250 aa had a ^D1D2D2^OPX architecture (Figure 4D; Figure S7).

We conclude that the coupling between ^D1D2^OPX proteins and OM β-barrel proteins is conserved and widespread in Gram-negative bacteria, and this coupling can be extended to include the ^D1D2D2^OPX variants.

### Conclusion

Based on the described experimental data and computational analyses, we propose that polysaccharides can be exported across the OM via a composite periplasmic OPX protein/OM β-barrel protein translocon (Figure 5). The domain architecture of the periplasmic OPX proteins can be ^D1D2^OPX or ^D1D2D2^OPX, giving rise to ^D1D2^OPX/β-barrel translocons and ^D1D2D2^OPX/β-barrel translocons. The ^D1D2^OPX/β-barrel translocons likely function with a PCP with extra-long periplasmic α-helical regions to span the periplasm and compensate for the short OPX protein (Figure 5). These systems are widespread in Gram-negative bacteria and add to the two established mechanisms for OM export, i.e. the Wza-like OPX protein in Wzx/Wzy-and ABC transporter-dependent pathways and the OM 16- to 18-stranded β-barrel protein in synthase-dependent pathways (Figure 5). In the case of the EpsX/EpsY proteins in *M. xanthus*, we have not shown that they interact directly. However, our model provides a framework for future biochemical and structural studies of EpsX and EpsY as well as of other polysaccharide biosynthesis pathways containing a ^D1D2^OPX or ^D1D2D2^OPX protein together with a coupled OM β-barrel protein.

**Figure 5.**
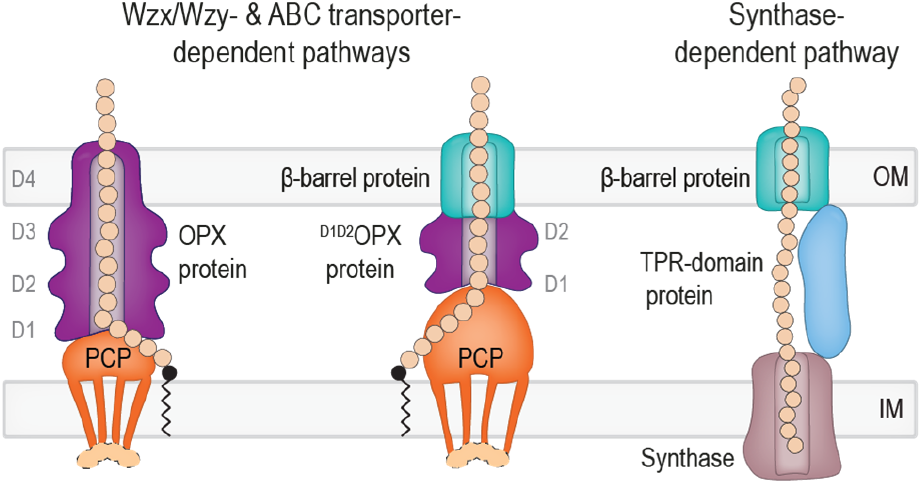
Three different mechanisms for polysaccharide export across the OM. Left schematic, in classical Wzx/Wzy- and ABC transporter-dependent pathways, polysaccharide transfer across the periplasm and OM is mediated by a complex composed of a PCP and an OPX protein with an α-helical barrel in the OM. Middle schematic, in pathways in which the OPX protein lacks domain D4 (^D1D2^OPX or ^D1D2D2^OPX proteins), a β- barrel protein constitutes the OM part of the composite OPX/β-barrel protein translocon. The D1-D4 and D1-D2 rings of the OPX proteins are indicated. In Wzx/Wzy-dependent pathways, the PCP can be associated to a cytoplasmic BY kinase (light orange). Synthesis, polymerization and translocation of the polysaccharide across the IM are not shown. Right schematic, in synthase-dependent pathways, transfer across the periplasm and the OM depends on a TPR domain-containing protein and a β-barrel protein translocon in the OM.

The existence of three systems for polysaccharide export across the OM poses the question of their evolutionary origin. Our current analysis does not suggest an evolutionary scenario for their emergence, and this will likely depend on detailed studies of the taxonomic distribution of the three systems.

## Acknowledgement

We thank Dr. Chris Whitfield for many helpful discussions and Dr. Dobromir Szadkowski for providing a Matlab script to illustrate AlphaFold model confidence plots. The Max Planck Society generously supported this work.

## Data Availability

The data that support the findings of this study are included in the manuscript or in the Supplementary Information. The source data underlying Figure 1A, 1D, 1E, 2B, 2C, 2E, 3B, 4A-C, 4D and Supplementary Figure S6B, S6C are provided as a Source Data file.

## Methods

### Strains and cell growth

All *M. xanthus* strains are derivatives of the WT DK1622 ^31^ and are listed in Table S1. Plasmids and oligonucleotides used are listed in Table S2 and Table S3. In-frame deletions were generated as described ^32^. The plasmid for the complementation experiment was integrated in a single copy by site specific recombination into the Mx8 *attB* site. All strains were verified by PCR. *M. xanthus* was grown at 32°C in 1% CTT broth (1% (w/v) Bacto Casitone, 10mM Tris-HCl pH 8.0, 1mM K_2_HPO_4_/KH_2_PO_4_ pH 7.6, 8mM MgSO_4_) or on 1.5% agar supplemented with 1% CTT, and kanamycin (50 μg mL^-1^) when required ^33^. Plasmids were propagated in *E. coli* Mach1 at 37°C in lysogeny broth (LB) ^34^ supplemented with kanamycin (50 μg mL^-1^).

### Detection of EPS

Colony-based EPS assays were performed as described ^17^. Briefly, exponentially growing cells were harvested (3 min, 6000× *g* at room temperature (RT)), resuspended in 1% CTT to a calculated density of 7×10^9^ cells ml^-1^, and 20μL aliquots placed on 0.5% agar plates supplemented with 0.5% CTT and 10 or 20μg mL^-1^ of Trypan blue or Congo red, respectively. Plates were incubated at 32°C and imaged at 24h.

### Motility assays

Motility assays were performed as described ^35^. Briefly, exponentially growing cells were harvested (3 min, 6000× *g*, RT) and resuspended in 1% CTT to a density of 7×10^9^ cells mL^-1^. 5μl of cell suspensions were placed on 0.5% and 1.5% agar (Invitrogen) supplemented with 0.5% CTT, and incubated at 32°C for 24h. Cells were imaged using a M205FA stereomicroscope (Leica) and a DMi8 inverted microscope (Leica) equipped with a Hamamatsu ORCA-flash V2 Digital CMOS camera (Hamamatsu Photonics) and DFC9000 GT camera (Leica), respectively.

### Proteomic analysis using data independent acquisition-mass spectrometry (DIA-MS)

The total proteome of *M. xanthus* cells grown on 1% CTT, 1.5% agar plates was determined as described ^24^. Peptide mixtures were then analyzed using liquid chromatography-mass spectrometry on an Exploris 480 instrument connected to an Ultimate 3000 RSLC nano with a Prowflow upgrade and a nanospray flex ion source (all Thermo Scientific). Peptide separation was performed on a reverse phase HPLC column (75μm × 42cm) packed in-house with C18 resin (2.4μm, Dr. Maisch). The following separating gradient was used: 94% solvent A (0.15% formic acid) and 6% solvent B (99.85% acetonitrile, 0.15% formic acid) to 25% solvent B over 95min and to 35% B for additional 25min at a flow rate of 300nL min^-1^. DIA-MS acquisition method was adapted from ^36^. Briefly, spray voltage was set to 2.0 kV, funnel RF level at 55, and heated capillary temperature at 275°C. For DIA experiments, full MS resolutions were set to 120.000 at m/z 200 with an Automatic Gain Control (AGC) target of 300% and max injection time (IT) of 50ms. Mass range was set to 350–1400. AGC target value for fragment spectra was set at 3000%. 49 windows of 15 Da were used with an overlap of 1 Da. Resolution was set to 15,000 and IT to 22ms. Stepped HCD collision energy of 25, 27.5 and 30% was used. MS1 data was acquired in profile, MS2 DIA data in centroid mode.

Analysis of DIA data was performed using DIA-NN version 1.8 ^37^ using a UniProt protein database for *M. xanthus*. Full tryptic digest was allowed with two missed cleavage sites, and oxidized methionines (variable) and carbamidomethylated cysteines (fixed). “Match between runs” and “remove likely interferences” options were enabled. The neural network classifier was set to the single-pass mode, and protein inference was based on “genes”. Quantification strategy was set to any LC (high accuracy). Cross-run normalization was set to “RT-dependent”. Library generation was set to “smart profiling”. The DIA-NN “report” output was used to sum the unique peptide intensities and identified proteins were filtered out if q-values exceeded 0.01. Protein intensities were normalized with the Cyclic Loess method using the R-package NormalyzerDE ^38^.

### RT-qPCR

Total RNA from *M. xanthus* cells grown on 1.5% agar supplemented with 1% CTwas extracted using the Monarch Total RNA Miniprep Kit (New England Biolabs (NEB)). Briefly, 10^9^ cells were scraped off the agar-plates and resuspended in 200μL lysis-buffer (100mM Tris-HCl pH 7.6, 1mg ml^-1^ lysozyme), and incubated at 25°C, 5min. RNA purification was performed according to manufacturer’s protocol. DNA was removed using Turbo DNase (Thermo Fisher Scientific) and DNase was removed by using the Monarch RNA Cleanup Kit (50μg; NEB). LunaScript RT SuperMix Kit (NEB) was used to generate cDNA using 1μg RNA. qPCR reactions were performed on an Applied Biosystems 7500 Real-Time PCR system using the Luna Universal qPCR MasterMix (NEB) with the primers listed in (Table S3). Data analysis was performed using the comparative *C*_T_ method ^39^. *MXAN_3298*, encoding the elongation factor Tu, served as internal reference gene as in ^25^.

### Bioinformatics

The phylogenetic tree of myxobacteria was prepared as in ^40^ in MEGA-X ^41^ using the Neighbor-Joining method ^42^ and the genome sequences listed in Table S4.

Full-length protein sequences or sequences in which the signal peptide was identified with SignalP 6.0 ^43^ and removed, were used for AlphaFold and AlphaFold-Multimer modeling via ColabFold ^44-46^ using the Alphafold2_mmseqs2 notebook with default settings, except recycles were set to six. Predicted Local Distance Difference Test (pLDDT) and predicted Alignment Error (pAE) graphs of the five models generated by Alphafold2_mmseqs2 notebook were made using a custom made Matlab R2020a (The MathWorks) script. Ranking of the models was performed based on combined pLDDT and pAE values, with the best ranked models used for further analysis and presentation. PyMOL version 2.4.1 (http://www.pymol.org/pymol) was used to analyze and visualize the models. Structural alignments were performed using the PyMOL Alignment plugin with default settings. Foldseek was used to identify protein homologs of EpsX in the PDB 100 database ^47^.

For the co-occurrence analysis of OPX and OM β-barrel proteins, we generated a database of 6,607 fully sequenced prokaryotic genomes that mirror the KEGG database ^48^ and comprehensively covers all major prokaryotic phyla. OPX proteins in these genomes were identified with HMMsearch of the HMMER package (v3.3) ^49^ using the Pfam ^50^ domain Poly_Export (PF02563) with the Pfam gathering threshold.

Subsequently, we generated a database corresponding to the proteins encoded by five genes up- and downstream of the identified OPX proteins. This database was searched for the Pfam-domains of polysaccharide synthesis proteins (ABC2_membrane (PF01061), ABC_tran (PF00005), Polysacc_synt (PF01943), Polysacc_synt3 (PF13440), Wzt_C (PF14524), Wzy_C (PF04932), Wzz (PF02706)) to (i) corroborate that the OPX protein is encoded in a polysaccharide synthesis gene cluster, and (ii) assign a pathway to the OPX protein. PCPs identified with the Wzz domain, were additionally analyzed for the presence of the tyrosine kinase-domain to assign them as PCP-2. OPX proteins that were not encoded in a polysaccharide synthesis gene cluster were excluded from further analysis. To remove redundancy in the dataset, OPX protein sequences were clustered to a 90% identity threshold using Cd-hit ^51^ and further analysis carried out on the resulting set of representative sequences.

To generate the phylogenetic tree of the OPX protein sequences, the sequence matching the Poly_export domain of the OPX proteins was aligned using MUSCLE ^52^ and a maximum likelihood tree inferred using IQ-Tree ^53^ with automated model selection and support calculated with 1000 ultrafast bootstrap replicates. The tree was visualized and annotated using iTOL ^54^.

The secondary structure of the OPX proteins was determined using S4pred ^55^, and then scanned for an α-helix within the last 20 C-terminal residues. C-terminal α-helices were considered positive, if they extended over more than nine residues, and did not contain more than one gap of two residues predicted to be non-helical.

β-barrel proteins were initially searched for by using hmmbuild to build an HMM from an alignment of 31 EpsX, ExoB and MXAN_1916 homologs generated with MUSCLE. In a second step, proteins from the KEGG orthology group VpsM (K20920) were aligned and used to build a second HMM. Both models were used to search the proteins encoded five genes up- and downstream of the genes encoding OPX proteins. β-barrel fold and OM localization of identified proteins was verified using PRED-TMBB2 ^56^ and PSORTb 3.0 ^57^. β- barrel proteins with less than 16 β-strands were classified as false-positive and were not considered further.

### Statistics

The Welch’s *t*-test was performed to determine the statistical differences between the samples.

## References

1 Morris, G. & Harding, S. in Encyclopedia of Microbiology 482–494 (Elsevier, 2009).

2 Sutherland, I. W. Novel and established applications of microbial polysaccharides. Trends Biotechnol. 16, 41–46 (1998).

3 Hall-Stoodley, L., Costerton, J. W. & Stoodley, P. Bacterial biofilms: from the natural environment to infectious diseases. Nat. Rev. Microbiol. 2, 95–108 (2004).

4 Low, K. E. & Howell, P. L. Gram-negative synthase-dependent exopolysaccharide biosynthetic machines. Curr. Opin. Struct. Biol. 53, 32–44 (2018).

5 Whitfield, C., Wear, S. S. & Sande, C. Assembly of bacterial capsular polysaccharides and exopolysaccharides. Annu. Rev. Microbiol. 74, 521–543 (2020).

6 Cuthbertson, L., Mainprize, I. L., Naismith, J. H. & Whitfield, C. Pivotal roles of the outer membrane polysaccharide export and polysaccharide copolymerase protein families in export of extracellular polysaccharides in gram-negative bacteria. Microbiol Mol Biol Rev 73, 155–177 (2009).

7 Dong, C. et al. Wza the translocon for E. coli capsular polysaccharides defines a new class of membrane protein. Nature 444, 226–229 (2006).

8 Nickerson, N. N. et al. Trapped translocation intermediates establish the route for export of capsular polysaccharides across Escherichia coli outer membranes. Proc Natl Acad Sci U S A 111, 8203–8208 (2014).

9 Reid, A. N. & Whitfield, C. Functional analysis of conserved gene products involved in assembly of Escherichia coli capsules and exopolysaccharides: Evidence for molecular recognition between Wza and Wzc for colanic acid biosynthesis. J. Bacteriol. 187, 5470–5481 (2005).

10 Collins, R. F. et al. The 3D structure of a periplasm-spanning platform required for assembly of group 1 capsular polysaccharides in Escherichia coli. Proc. Natl. Acad. Sci USA 104, 2390–2395 (2007).

11 Whitney, J. C. et al. Structural basis for alginate secretion across the bacterial outer membrane. Proc. Natl. Acad. Sci USA 108, 13083–13088 (2011).

12 Wang, Y. et al. Structural basis for translocation of a biofilm-supporting exopolysaccharide across the bacterial outer membrane. J. Biol. Chem. 291, 10046–10057 (2016).

13 Acheson, J. F., Derewenda, Z. S. & Zimmer, J. Architecture of the cellulose synthase outer membrane channel and its association with the periplasmic TPR domain. Structure 27, 1855-1861.e1853 (2019).

14 Pérez-Burgos, M. & Søgaard-Andersen, L. Biosynthesis and function of cell-surface polysaccharides in the social bacterium Myxococcus xanthus. Biol. Chem. 401, 1375–1387 (2020).

15 Holkenbrink, C., Hoiczyk, E., Kahnt, J. & Higgs, P. I. Synthesis and assembly of a novel glycan layer in Myxococcus xanthus spores. J. Biol. Chem. 289, 32364–32378 (2014).

16 Kahnt, J. et al. Profiling the outer membrane proteome during growth and development of the social bacterium Myxococcus xanthus by selective biotinylation and analyses of outer membrane vesicles. J. Proteome Res. 9, 5197–5208 (2010).

17 Pérez-Burgos, M. et al. Characterization of the exopolysaccharide biosynthesis pathway in Myxococcus xanthus. J. Bacteriol. 202, e00335–00320 (2020).

18 Islam, S. T. et al. Modulation of bacterial multicellularity via spatio-specific polysaccharide secretion. PLOS Biol. 18, e3000728 (2020).

19 Lu, A. et al. Exopolysaccharide biosynthesis genes required for social motility in Myxococcus xanthus. Mol. Microbiol. 55, 206–220 (2005).

20 Rahn, A., Beis, K., Naismith, J. H. & Whitfield, C. A novel outer membrane protein, Wzi, is involved in surface assembly of the Escherichia coli K30 group 1 capsule. J Bacteriol 185, 5882–5890 (2003).

21 Bushell, S. R. et al. Wzi is an outer membrane lectin that underpins group 1 capsule assembly in Escherichia coli. Structure 21, 844–853 (2013).

22 Jakobczak, B., Keilberg, D., Wuichet, K. & Søgaard-Andersen, L. Contact-and protein transfer-dependent stimulation of assembly of the gliding motility machinery in Myxococcus xanthus. PLOS Genet 11, e1005341 (2015).

23 Friedrich, C., Bulyha, I. & Søgaard-Andersen, L. Outside-in assembly pathway of the type IV pili system in Myxococcus xanthus. J Bacteriol 196, 378–390 (2014).

24 Treuner-Lange, A. et al. PilY1 and minor pilins form a complex priming the type IVa pilus in Myxococcus xanthus. Nat Commun 11, 5054 (2020).

25 Herfurth, M. et al. A noncanonical cytochrome c stimulates calcium binding by PilY1 for type IVa pili formation. Proc. Natl. Acad. Sci USA 119, e2115061119 (2022).

26 Sande, C. et al. Structural and functional variation in outer membrane polysaccharide export (OPX) oroteins from the two major capsule assembly pathways present in Escherichia coli. J Bacteriol 201 (2019).

27 Sathiyamoorthy, K., Mills, E., Franzmann, T. M., Rosenshine, I. & Saper, M. A. The crystal structure of Escherichia coli Group 4 capsule protein GfcC reveals a domain organization resembling that of Wza. Biochemistry 50, 5465–5476 (2011).

28 Nesper, J. et al. Translocation of Group 1 capsular polysaccharide in Escherichia coli serotype K30: Structural and functional analysis of the outer membrane lipoprotein Wza. J. Biol. Chem. 278, 49763–49772 (2003).

29 Yang, Y. et al. The molecular basis of regulation of bacterial capsule assembly by Wzc. Nat Commun 12, 4349 (2021).

30 Fong, J. C. N., Syed, K. A., Klose, K. E. & Yildiz, F. H. Role of Vibrio polysaccharide (vps) genes in VPS production, biofilm formation and Vibrio cholerae pathogenesis. Microbiology 156, 2757–2769 (2010).

31 Kaiser, D. Social gliding is correlated with the presence of pili in Myxococcus xanthus. Proc. Natl. Acad. Sci. USA 76, 5952–5956 (1979).

32 Shi, X. et al. Bioinformatics and experimental analysis of proteins of two-component systems in Myxococcus xanthus. J Bacteriol 190, 613–624 (2008).

33 Hodgkin, J. & Kaiser, D. Cell-to-cell stimulation of movement in nonmotile mutants of Myxococcus. Proc Natl Acad Sci U S A 74, 2938–2942 (1977).

34 Sambrook, J. & Russell, D. W. Molecular Cloning: A Laboratory Manual. 3rd edn, (Cold Spring Harbor Laboratory Press), (2001).

35 Shi, W. & Zusman, D. R. The two motility systems of Myxococcus xanthus show different selective advantages on various surfaces. Proc Natl Acad Sci U S A 90, 3378–3382 (1993).

36 Bekker-Jensen, D. B. et al. A compact Quadrupole-Orbitrap Mass Spectrometer with FAIMS interface improves proteome coverage in short LC Gradients. Mol Cell Proteomics 19, 716–729 (2020).

37 Demichev, V., Messner, C. B., Vernardis, S. I., Lilley, K. S. & Ralser, M. DIA-NN: neural networks and interference correction enable deep proteome coverage in high throughput. Nat. Methods 17, 41–44 (2020).

38 Willforss, J., Chawade, A. & Levander, F. NormalyzerDE: Online tool for improved normalization of omics expression data and high-sensitivity differential expression analysis. J. Proteome Res. 18, 732–740 (2019).

39 Schmittgen, T. D. & Livak, K. J. Analyzing real-time PCR data by the comparative C(T) method. Nat Protoc 3, 1101–1108 (2008).

40 Pérez-Burgos, M., García-Romero, I., Valvano, M. A. & Søgaard Andersen, L. Identification of the Wzx flippase, Wzy polymerase and sugar-modifying enzymes for spore coat polysaccharide biosynthesis in Myxococcus xanthus. Mol. Microbiol. 113, 1189–1208 (2020).

41 Kumar, S., Stecher, G. & Tamura, K. MEGA7: Molecular evolutionary genetics analysis version 7.0 for bigger datasets. Mol Biol Evol 33, 1870–1874 (2016).

42 Saitou, N. & Nei, M. The Neighbor-Joining Method -a new method for reconstructing phylogenetic trees. Mol Biol Evol 4, 406–425 (1987).

43 Teufel, F. et al. SignalP 6.0 predicts all five types of signal peptides using protein language models. Nat Biotechnol, https://doi.org/10.1038/s41587-41021-01156-41583 (2022).

44 Jumper, J. et al. Highly accurate protein structure prediction with AlphaFold. Nature 596, 583–589 (2021).

45 Mirdita M., S. K., Moriwaki Y., Heo L., Ovchinnikov S., Steinegger M. ColabFold -Making protein folding accessible to all. bioRxiv, 10.1101/2021.1108.1115.456425 (2021).

46 Evans R., O. N. M., Pritzel A., Antropova N., Senior A., Green T., Žídek A., Bates R., Blackwell S., Yim J., Ronneberger O., Bodenstein S., Zielinski M., Bridgland A., Potapenko A., Cowie A., Tunyasuvunakool K., Jain R., Clancy E., Kohli P., Jumper J., Hassabis D. Protein complex prediction with AlphaFold-Multimer. bioRxiv, 10.1101/2021.1110.1104.463034 (2021).

47 van Kempen, M. et al. Foldseek: fast and accurate protein structure search. bioRxiv, 10.1101/2022.1102.1107.479398 (2021).

48 Kanehisa, M. & Goto, S. KEGG: kyoto encyclopedia of genes and genomes. Nucleic Acids Res 28, 27–30 (2000).

49 Eddy, S. R. Accelerated profile HMM searches. PLOS Comput Biol 7, e1002195 (2011).

50 Mistry, J. et al. Pfam: The protein families database in 2021. Nucl. Acids Res. 49, D412–D419 (2021).

51 Li, W. & Godzik, A. Cd-hit: a fast program for clustering and comparing large sets of protein or nucleotide sequences. Bioinformatics 22, 1658–1659 (2006).

52 Edgar, R. C. MUSCLE: multiple sequence alignment with high accuracy and high throughput. Nucl Acids Res 32, 1792–1797 (2004).

53 Minh, B. Q. et al. IQ-TREE 2: New models and efficient methods for phylogenetic inference in the genomic era. Mol Biol Evol 37, 1530–1534 (2020).

54 Letunic, I. & Bork, P. Interactive Tree Of Life (iTOL) v5: an online tool for phylogenetic tree display and annotation. Nucl Acids Res 49, W293–W296 (2021).

55 Moffat, L. & Jones, D. T. Increasing the accuracy of single sequence prediction methods using a deep semi-supervised learning framework. Bioinformatics (2021).

56 Tsirigos, K. D., Elofsson, A. & Bagos, P. G. PRED-TMBB2: improved topology prediction and detection of beta-barrel outer membrane proteins. Bioinformatics 32, i665–i671 (2016).

57 Yu, N. Y. et al. PSORTb 3.0: improved protein subcellular localization prediction with refined localization subcategories and predictive capabilities for all prokaryotes. Bioinformatics 26, 1608–1615 (2010).

58 Schulz, G. E. Beta-barrel membrane proteins. Curr Opin Struct Biol 10, 443–447 (2000).

59 Wu, S. S. & Kaiser, D. Regulation of expression of the pilA gene in Myxococcus xanthus. J Bacteriol 179, 7748–7758 (1997).

