## Supplementary Information for "Evidence for a widespread third system for bacterial polysaccharide export across the outer membrane comprising a composite OPX/β-barrel translocon"

**This file contains:**

- Supplementary Figures 1-7
- Supplementary Experimental Procedures
- Supplementary Tables 1-4
- Supplementary References

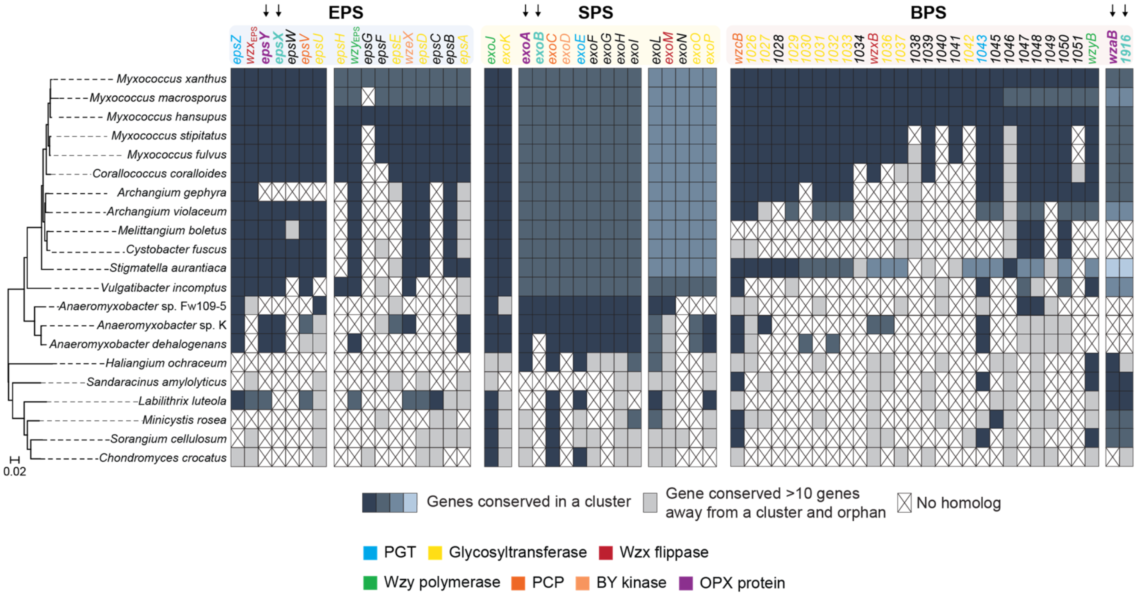


**Figure S1. The gene clusters for EPS, SPS and BPS biosynthesis in *M. xanthus* are conserved in other myxobacteria.**

Left, 16S rRNA-based tree of fully sequenced myxobacteria. Right, a reciprocal best BLASTP hit method was used to identify orthologs as described [3,6]. 10 genes was considered the maximum distance for a gene to be in a cluster. For each pathway, genes in the same cluster are marked in identical colors. Genes within a distance <10 genes were considered as part of the same cluster. Clusters were considered separate when they were more than 10 genes. Conserved but orphan genes are colored in light grey. Genes without an ortholog are indicated by a cross. In the top row, *M. xanthus* genes are color-coded according to the code shown below. Black arrows indicate synteny of genes encoding the OPX protein (purple) and genes encoding an 18-stranded β-barrel protein (teal) conserved in *eps*, *sps* and *bps* gene clusters in myxobacteria. In the *eps* gene cluster, WzeX is important for EPS synthesis and was proposed to act as the BY-kinase partner of EpsV [4,5], the serine O-acetyltransferase EpsC is thought to be involved in sugar nucleotide precursor biosynthesis and is not important for EPS biosynthesis [1,3,5], EpsB is a predicted glycoside hydrolase, which is not important for EPS synthesis [1]. EpsW is a response regulator involved in regulation of EPS synthesis [7]. *epsG* and *epsF* are annotated as encoding a magnesium transporter and a hybrid response regulator/ histidine kinase, respectively, and *epsF* is not required for EPS synthesis [1]. In the *bps* cluster, *wzcB* encodes a PCP protein with an N-terminal BY kinase domain. For a more detailed description of proteins encoded by the *sps* and *bps* gene clusters and not described in the color code, see [3,5,6].

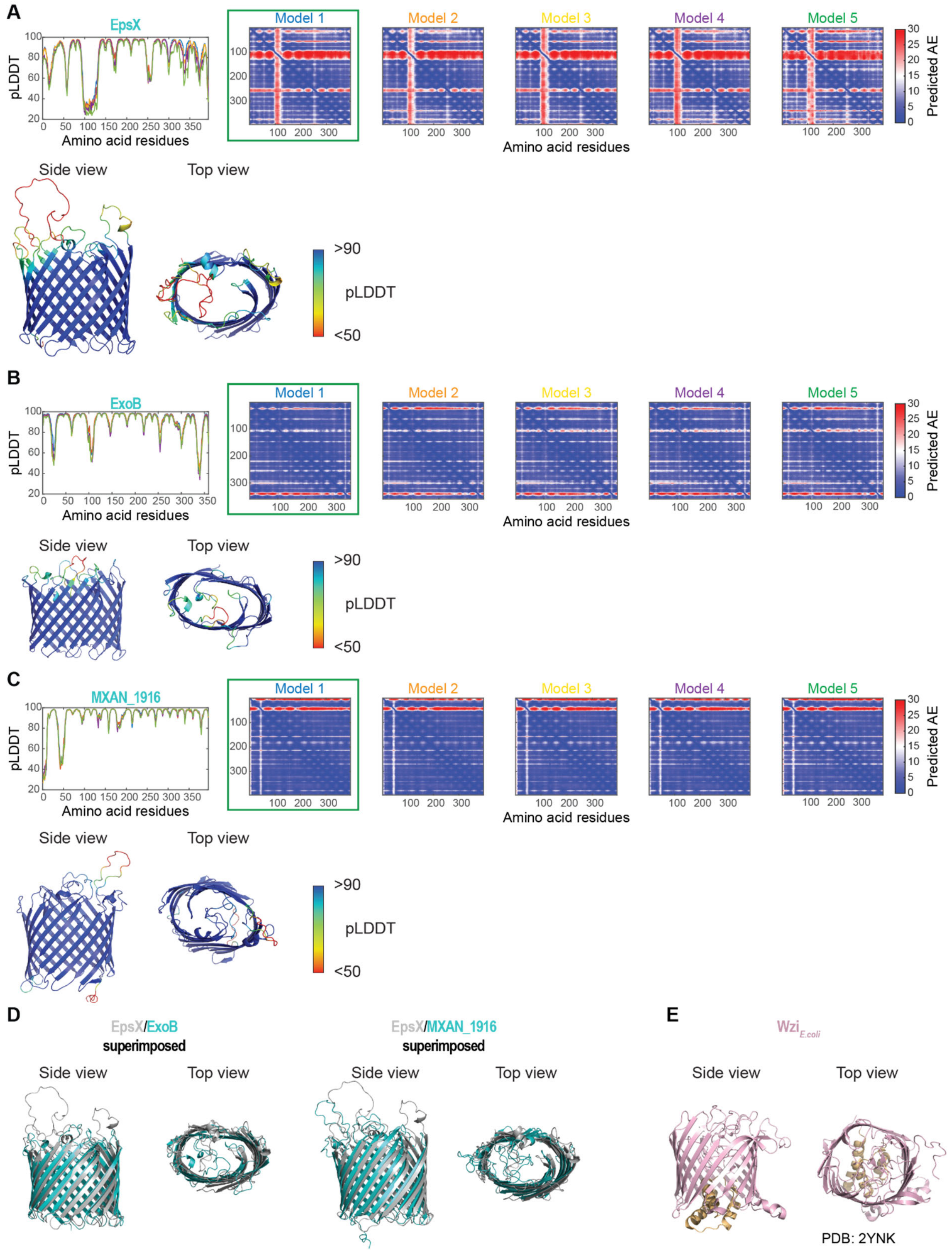


**Figure S2. AlphaFold models of EpsX, ExoB and MXAN_1916.**
**A, B, C.** pLDDT (Predicted Local Distance Difference Test) and pAE (predicted Alignment Error) plots for five models of EpsX (A), ExoB (B) and MXAN_1916 (C) as predicted by AlphaFold. For all three proteins, model rank 1 (marked by green box) was used for further analysis and is shown colored based on pLDDT. Mobility and low sequence conservation are typical features of the extracellular loops of β-barrels [8], likely explaining the low spatial

confidence in loop positioning in the EpsX, ExoB and MXAN_1916 models. Signal peptides were removed before generating a model.

**D.** Superimposition of AlphaFold predicted structures of EpsX and ExoB, and EpsX and MXAN_1916. Left panel, EpsX is colored in grey and ExoB in teal. ExoB aligns to EpsX with an RMSD of 3.427Å over 232 $C_{\alpha}$. Right panel, EpsX is colored in grey and MXAN_1916 in teal. MXAN_1916 aligns to EpsX with an RMSD of 4.484Å over 320 $C_{\alpha}$.

**E**. The solved structure of Wzi (PDB 2YNK) [9]. The protein is colored in light pink, except for the N-terminal α-helical bundle that closes the barrel to the periplasm [9], which is colored in light orange.

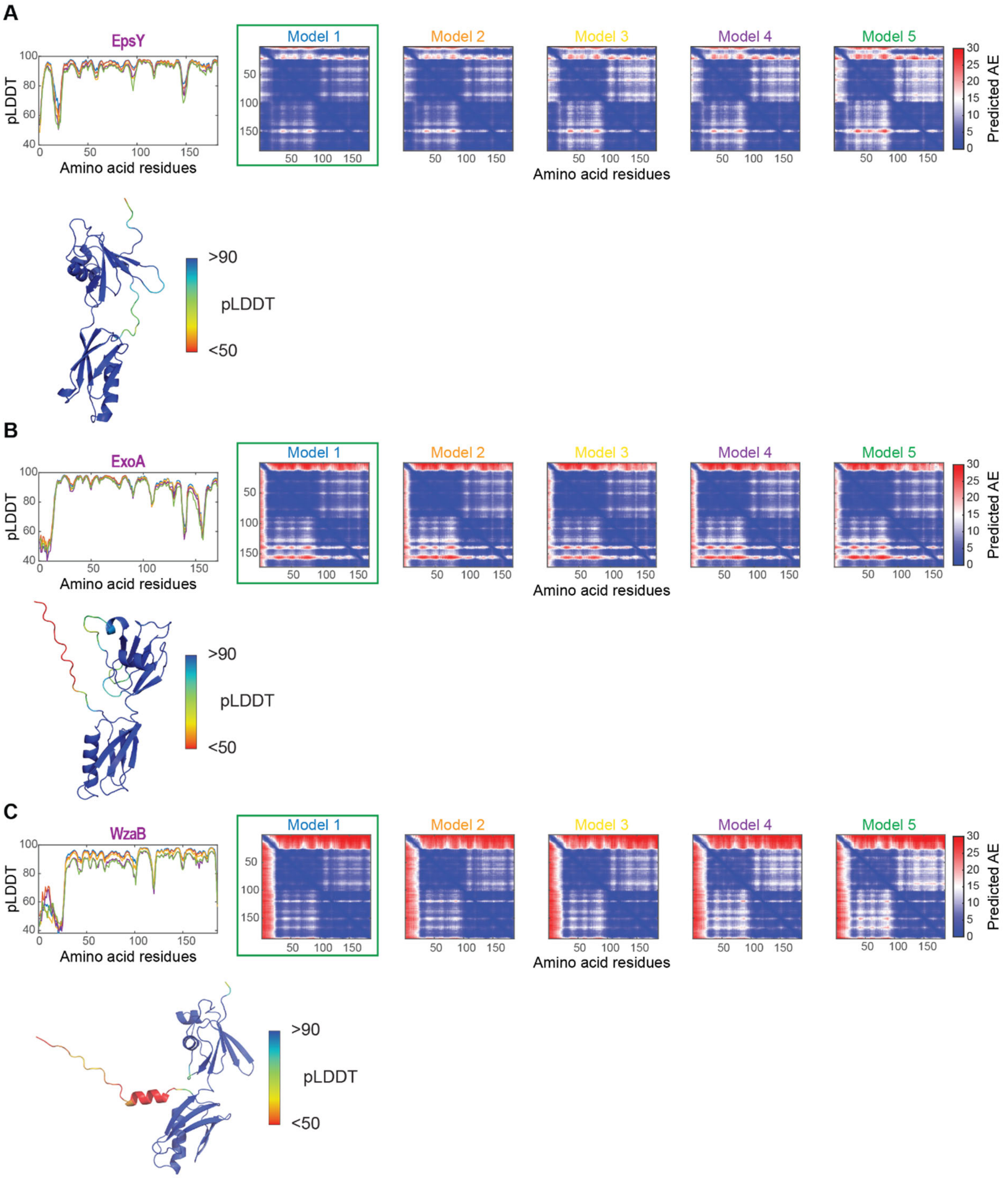


**Figure S3. AlphaFold models of EpsY, ExoA and WzaB.**
**A, B, C.** pLDDT and pAE plots for five models of the indicated proteins as predicted by AlphaFold. For all three proteins, model rank 1 (marked by green box) was used for further analysis and is shown colored based on pLDDT. Signal peptides were removed before generating a model.

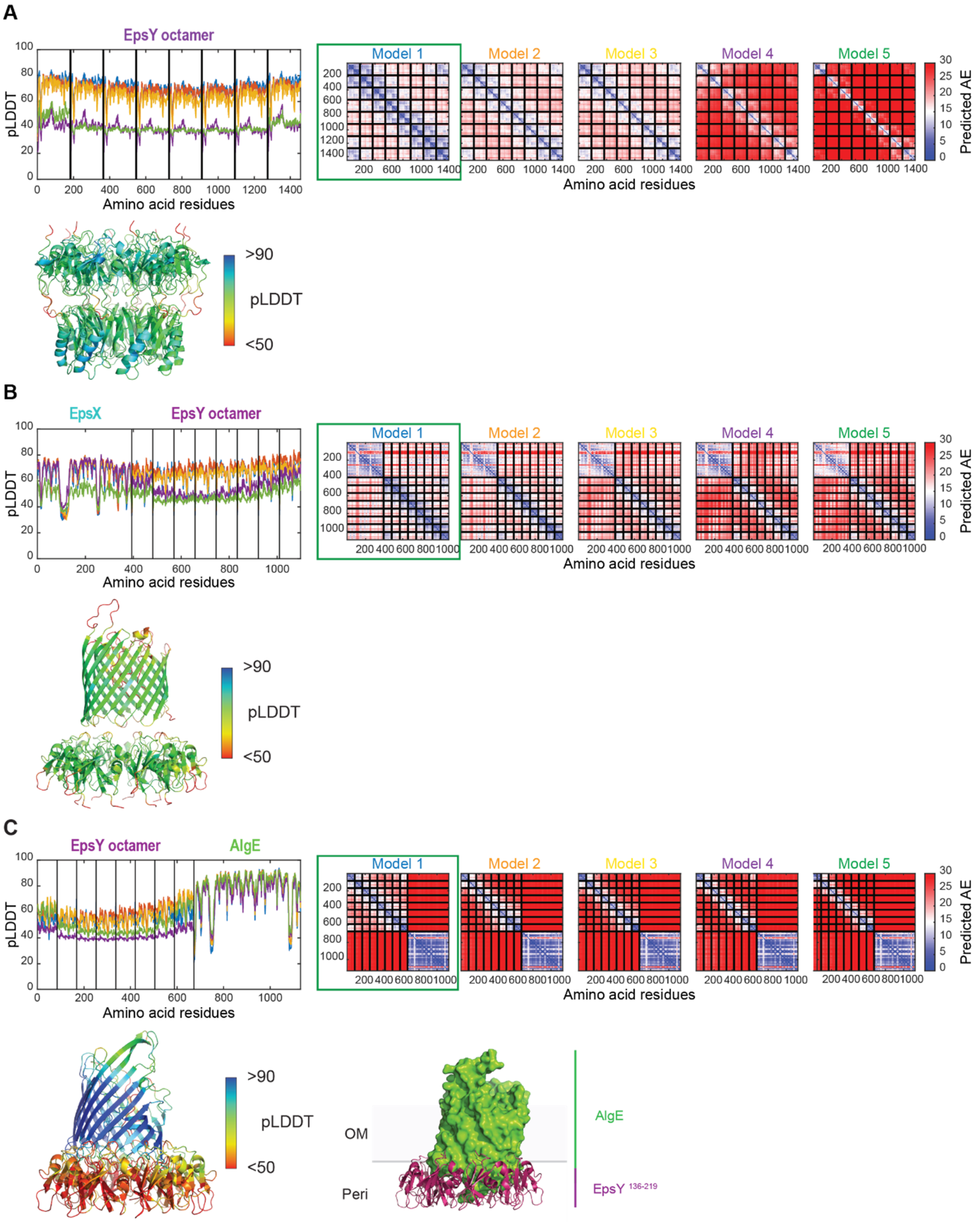


**Figure S4. AlphaFold-Multimer models of EpsY octamer, EpsX/EpsY$^{136\text{-}219}$ and AlgE/EpsY$^{136\text{-}219}$**

**A, B, C.** pLDDT and pAE plots for five models of the indicated protein complexes as predicted by AlphaFold-Multimer. For all three complexes, model rank 1 (marked by green box) was used for further analysis and is shown colored based on pLDDT. In **C**, the AlgE/EpsY$^{136\text{-}219}$ heterocomplex is also shown with AlgE surface-rendered. Signal peptides were removed before generating a model.

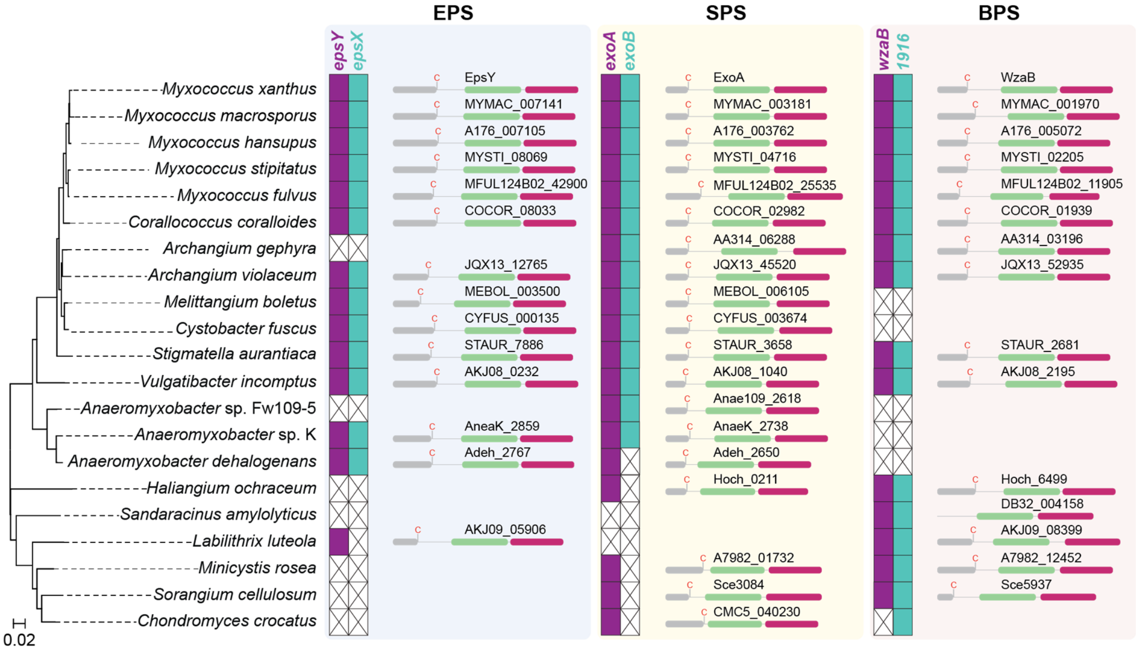


**Figure S5. Conservation of domain structure of $^{D1D2}$OPX proteins in myxobacteria.** Excerpt from Figure S1 focusing on the synteny of $^{D1D2}$OPX protein encoding genes (purple) and OM 18-stranded β-barrel protein encoding genes (teal) in the *eps*, *sps* and *bps* gene clusters in myxobacterial genomes. In the right panel of each pathway, the domain architecture of the $^{D1D2}$OPX proteins is colored as in Figure 2A-B.

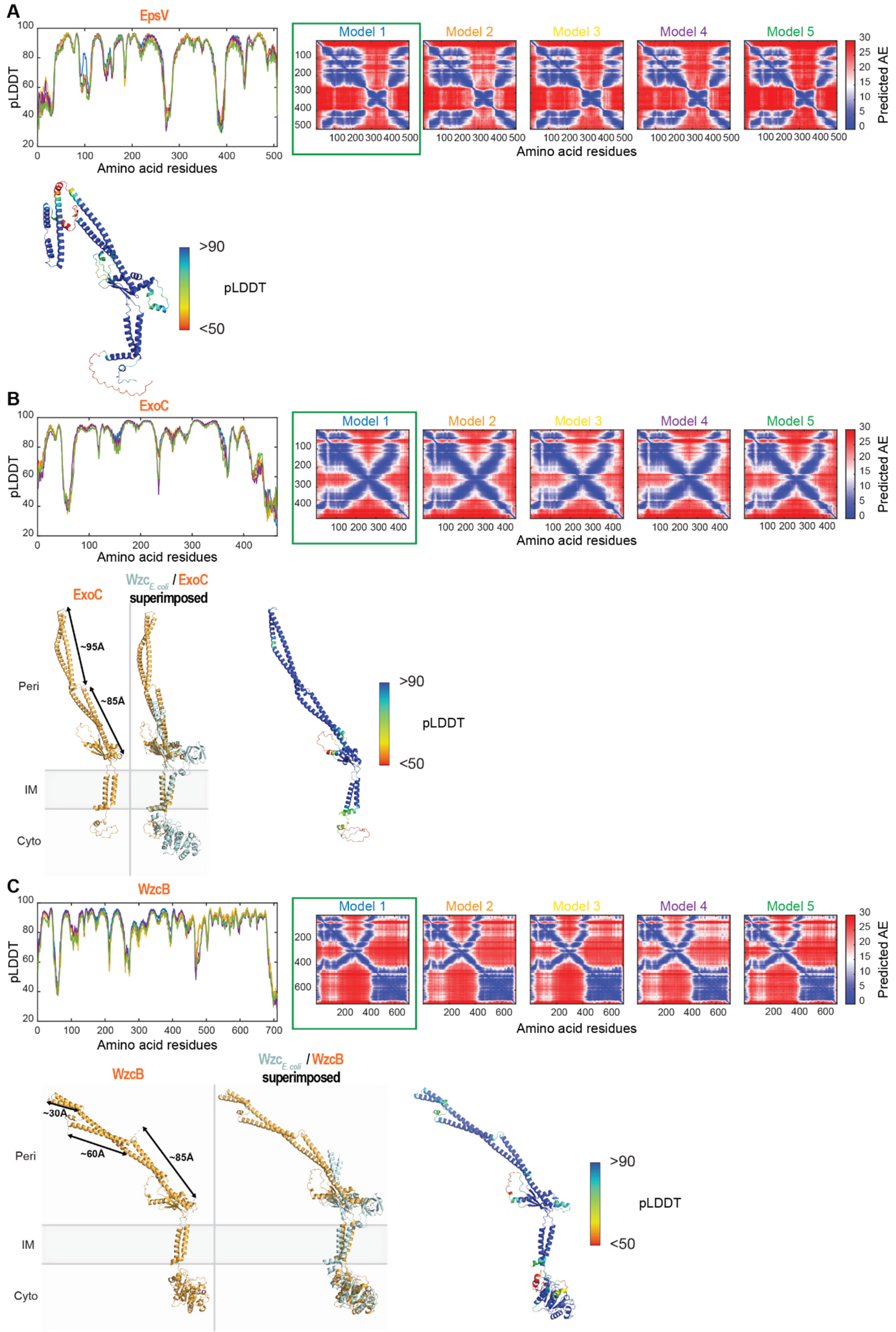


**Figure S6. AlphaFold models of EpsV, ExoC and WzcB.**

**A, B, C.** pLDDT and pAE plots for five models of the indicated proteins as predicted by AlphaFold. For all three proteins, model rank 1 (marked by green box) was used for further analysis and is shown colored based on pLDDT. In the lower left panels in **B** and **C**, the proteins are shown in orange and arrows indicate the length of the extended α-helical stretches. Middle panels, superimposition of a protomer from the solved octameric structure of Wzc (PDB 7NHR) [10] and the relevant protein model. ExoC aligns to Wzc with an RMSD of 2.706Å over 776 $C_\alpha$ and WzcB aligns to Wzc with an RMSD of 3.946Å over 1719 $C_\alpha$. Source data for **B, C** are provided in the Source Data file.

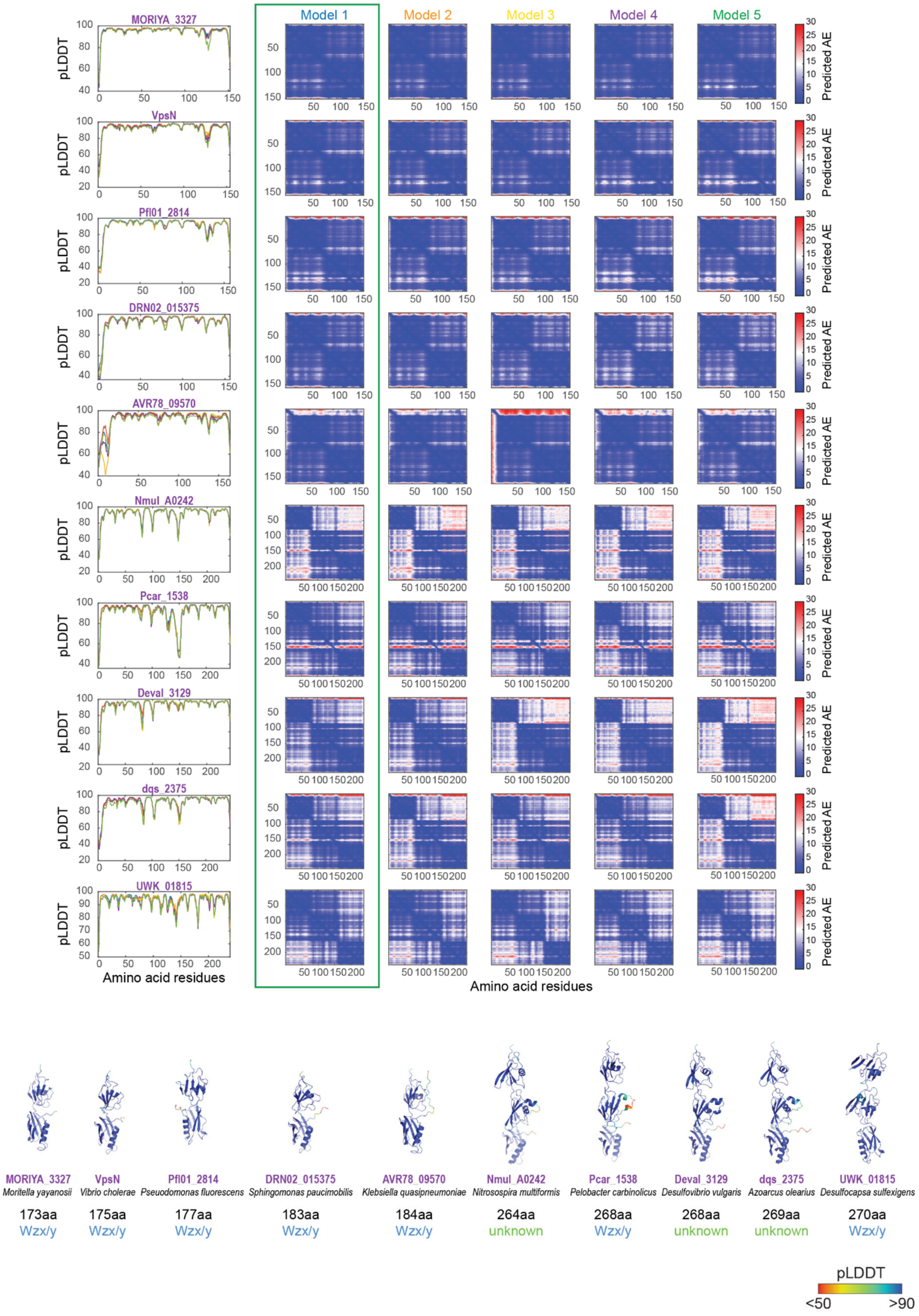


**Figure S7. AlphaFold models of the indicated proteins.**
pLDDT and pAE plots for five models of the indicated proteins predicted by AlphaFold. For all 10 proteins, model rank 1 (marked by green box) was used for further analysis and is

shown colored based on pLDDT below. Signal peptides were removed before generating a model.

### Supplementary Methods

Plasmid construction. All oligonucleotides used are listed in Table S3. All constructed plasmids were verified by DNA sequencing.

**pJSc004** (for generation of in-frame deletion of *epsX*): up- and downstream fragments were amplified from genomic DNA of DK1622 using the primer pairs 7418-A/7418-B and 7418-C/7418-D, respectively. Subsequently, the AB and CD fragments were used as templates for an overlapping PCR with the primer pair 7418-A/7418-D to generate the AD fragment. The AD fragment was digested with KpnI/XbaI and cloned in pBJ114.

**pJSc007** (for generation of a complementation strain ectopically expressing *epsX* from the *pilA* promotor): *MXAN_7418* was amplified from genomic DNA of DK1622 using 7418-P*pilA*-for and 7418-Pnat/P*pilA*-rev. Subsequently, the PCR fragment was digested with XbaI/HindIII, and cloned into pSW105.

**Table S1.** Strains used in this work

| **Strain** | **Genotype** | **Reference** |
|---|---|---|
| ***M. xanthus*** | | |
| DK1622 | WT | [11] |
| DK10410 | Δ*pilA* | [12] |
| SA3922 | Δ*gltB* | [13] |
| SA7400 | Δ*epsZ* | [3] |
| SA7406 | Δ*epsV* | [3] |
| SA7408 | Δ*epsY* | [3] |
| SA11550 | Δ*epsX* | This study |
| SA11554 | Δ*epsX*/$P_{pilA}$ *epsX* | This study |
| ***E. coli*** | | |
| Mach1 | Δ*recA*1398 *endA*1 *tonA* Φ80Δ*lacM*15 Δ*lacX*74 *hsdR*($r_K^-$ $m_K^+$) | Invitrogen |

**Table S2.** Plasmids used in this work

| Plasmid | Description | Reference |
|---|---|---|
| pBJ114 | Km$^r$, *galK* | [14] |
| pSW105 | Km$^r$, P*pilA* | [15] |
| pJSc004 | pBJ114, in-frame deletion construct for *epsX* (*MXAN_7418*), Km$^r$ | This study |
| pJSc007 | pSW105, complementation construct for *epsX* (*MXAN_7418*) expressed from the *pilA* promoter, Km$^r$ | This study |

**Table S3.** Oligonucleotides used in this work[1]

| Primer name | Sequence 5'-3' | Brief descrption |
|---|---|---|
| 7418-A | TTTGGTACCGGGTGCGCATCACCGTGG | For Δ*epsX* |
| 7418-B | CCGCACGGACGCGACGGTGAGGACCGT | For Δ*epsX* |
| 7418-C | ACCGTCGCGTCCGTGCGGCAAACTGGT | For Δ*epsX* |
| 7418-D | TTTTCTAGAAGGAACCAGTGCCGCAGC | For Δ*epsX* |
| 7418-E | ATGCTTTCGGCGCTGGGC | For Δ*epsX* |
| 7418-F | GTTGCGCTGCGTCAGCAT | For Δ*epsX* |
| 7418-G | ACACCGACGTCACCCCGC | For Δ*epsX* |
| 7418-H | GGATGCTGTCCCCACGAC | For Δ*epsX* |
| 7418-P*pilA*-for | AAATCTAGAGTGCTGGGGACGGTCCTCA | For complementation of Δ*epsX* |
| 7418-Pnat/P*pilA*-rev | CCCAAGCTTTCAGAGAATACCAGTTTGCCG | For complementation of Δ*epsX* |
| 7417-q-for-2 | AGGACTACATCAACCACCCC | RT-qPCR for *epsY* |
| 7417-q-rev-2 | TGACGAAGATGCGGTCCTTG | RT-qPCR for *epsY* |
| 7418-q-for-5 | CTCCTGGGCCTGGAAATTCG | RT-qPCR for *epsX* |
| 7418-q-rev-5 | CATGTGCTGGATTTCGGTGC | RT-qPCR for *epsX* |
| 7421-q-for-2 | CGACGCGGTCTTCTTTTTGA | RT-qPCR for *epsV* |
| 7421-q-rev-2 | CATGATTTTGCTGACGCCCA | RT-qPCR for *epsV* |

[1] Underlined sequences indicate restriction sites.

**Table S4.** Fully sequenced myxobacterial genomes used for the 16S RNA tree

| Species and strain name |
|---|
| *Anaeromyxobacter dehalogenans* 2CP-C |
| *Anaeromyxobacter* sp. Fw109-5 |
| *Anaeromyxobacter* sp. K |
| *Archangium gephyra* DSM 2261 |
| *Archangium violaceum* Cb SDU34 |
| *Chondromyces crocatus* Cm c5 |
| *Corallococcus coralloides* DSM 2259 |
| *Cystobacter fuscus* DSM 52655 |
| *Haliangium ochraceum* DSM 14365 |
| *Labilithrix luteola* DSM 27648 |
| *Melittangium boletus* DSM 14713SG |
| *Minicystis rosea* DSM 24000 |
| *Myxococcus macrosporus* DSM 14675 |
| *Myxococcus hansupus* (*Myxococcus* sp. mixupus) |
| *Myxococcus stipitatus* DSM 14675 |
| *Myxococcus xanthus* DK1622 |
| *Sandaracinus amylolyticus* DSM 53668 |
| *Sorangium cellulosum* So ce 56 |
| *Stigmatella aurantiaca* DW4/3-1 |
| *Vulgatibacter incomptus* DSM 27710 |

### Supplementary References


1 Lu, A. *et al.* Exopolysaccharide biosynthesis genes required for social motility in *Myxococcus xanthus*. *Mol. Microbiol.* **55**, 206-220 (2005).

2 Youderian, P. & Hartzell, P. L. Transposon insertions of *magellan-4* that impair social gliding motility in *Myxococcus xanthus. Genetics* **172**, 1397-1410 (2006).

3 Pérez-Burgos, M. *et al.* Characterization of the exopolysaccharide biosynthesis pathway in *Myxococcus xanthus*. *J. Bacteriol.* **202**, e00335-00320 (2020).

4 Pérez-Burgos, M. & Søgaard-Andersen, L. Biosynthesis and function of cell-surface polysaccharides in the social bacterium *Myxococcus xanthus*. *Biol. Chem.* **401**, 1375-1387 (2020).

5 Islam, S. T. *et al.* Modulation of bacterial multicellularity via spatio-specific polysaccharide secretion. *PLOS Biol.* **18**, e3000728 (2020).

6 Pérez-Burgos, M., García-Romero, I., Valvano, M. A. & Søgaard Andersen, L. Identification of the Wzx flippase, Wzy polymerase and sugar-modifying enzymes for spore coat polysaccharide biosynthesis in *Myxococcus xanthus*. *Mol. Microbiol.* **113**, 1189-1208 (2020).

7 Black, W. P., Wang, L., Davis, M. Y. & Yang, Z. The orphan response regulator EpsW is a substrate of the DifE kinase and it regulates exopolysaccharide in *Myxococcus xanthus*. *Sci. Rep.* **5**, 17831 (2015).

8 Schulz, G. E. Beta-barrel membrane proteins. *Curr Opin Struct Biol* **10**, 443-447 (2000).

9 Bushell, S. R. *et al.* Wzi is an outer membrane lectin that underpins group 1 capsule assembly in *Escherichia coli*. *Structure* **21**, 844-853 (2013).

10 Yang, Y. *et al.* The molecular basis of regulation of bacterial capsule assembly by Wzc. *Nat Commun* **12**, 4349 (2021).

11 Kaiser, D. Social gliding is correlated with the presence of pili in *Myxococcus xanthus*. *Proc. Natl. Acad. Sci. USA* **76**, 5952-5956 (1979).

12 Wu, S. S. & Kaiser, D. Regulation of expression of the *pilA* gene in *Myxococcus xanthus*. *J Bacteriol* **179**, 7748-7758 (1997).

13 Jakobczak, B., Keilberg, D., Wuichet, K. & Søgaard-Andersen, L. Contact- and protein transfer-dependent stimulation of assembly of the gliding motility machinery in *Myxococcus xanthus*. *PLOS Genet* **11**, e1005341 (2015).

14 Julien, B., Kaiser, A. D. & Garza, A. Spatial control of cell differentiation in *Myxococcus xanthus*. *Proc Natl Acad Sci U S A* **97**, 9098-9103 (2000).

15 Jakovljevic, V., Leonardy, S., Hoppert, M. & Søgaard-Andersen, L. PilB and PilT are ATPases acting antagonistically in type IV pilus function in *Myxococcus xanthus*. *J Bacteriol* **190**, 2411-2421 (2008).
